# Genetic-epigenetic tissue mapping for plasma DNA: applications in prenatal testing, transplantation and oncology

**DOI:** 10.1101/2020.11.13.381178

**Authors:** Wanxia Gai, Ze Zhou, Sean Agbor-Enoh, Xiaodan Fan, Sheng Lian, Peiyong Jiang, Suk Hang Cheng, John Wong, Stephen L. Chan, Moon Kyoo Jang, Yanqin Yang, Raymond H. S. Liang, Wai Kong Chan, Edmond S. K. Ma, Tak Y. Leung, Rossa W. K. Chiu, Hannah Valantine, K. C. Allen Chan, Y. M. Dennis Lo

**Affiliations:** Li Ka Shing Institute of Health Sciences, The Chinese University of Hong Kong, Shatin, New Territories, Hong Kong SAR, China; Department of Chemical Pathology, The Chinese University of Hong Kong, Prince of Wales Hospital, Shatin, New Territories, Hong Kong SAR, China; Genomic Research Alliance for Transplantation (GRAfT), 10 Center Drive, 7S261, Bethesda, MD 20982, United States; Division of Pulmonary and Critical Care Medicine, The Johns Hopkins School of Medicine, 1830 East Monument Street, Baltimore, MD, United States; Division of Intramural Research, National Heart, Lung and Blood Institute, 10 Center Drive, 7S261, Bethesda, MD 20982, United States; Department of Statistics, The Chinese University of Hong Kong, Shatin, New Territories, Hong Kong SAR, China; Department of Surgery, The Chinese University of Hong Kong, Prince of Wales Hospital, Shatin, New Territories, Hong Kong SAR, China; Department of Clinical Oncology, The Chinese University of Hong Kong, Prince of Wales Hospital, Shatin, New Territories, Hong Kong SAR, China; Comprehensive Oncology Centre, Hong Kong Sanatorium & Hospital, Hong Kong SAR, China; Department of Pathology, Hong Kong Sanatorium & Hospital, Hong Kong SAR, China; Department of Obstetrics and Gynaecology, The Chinese University of Hong Kong, Prince of Wales Hospital, Shatin, New Territories, Hong Kong SAR, China

**Author notes:** **Corresponding Author:** Y.M. Dennis Lo, The Chinese University of Hong Kong, Prince of Wales Hospital, 30–32 Ngan Shing Street, Shatin, New Territories, Hong Kong SAR, China. **Note:** W. Gai, and Z. Zhou contributed equally to this work.

## Abstract

We developed <u>G</u>enetic-<u>E</u>pigenetic <u>T</u>issue <u>Map</u>ping (GETMap) to determine the tissue composition of plasma DNA carrying genetic variants not present in the constitutional genome through comparing their methylation profiles with relevant tissues. We validated this approach by showing that, in pregnant women, circulating DNA carrying fetal-specific alleles was entirely placenta-derived. In lung-transplant recipients, we showed that, at 72 hours after transplantation, the lung contributed only a median of 17% to the plasma DNA carrying donor-specific alleles and hematopoietic cells contributed a median of 78%. In hepatocellular cancer patients, the liver was identified as the predominant source of plasma DNA carrying tumor-specific mutations. In a pregnant woman with lymphoma, plasma DNA molecules carrying cancer mutations and fetal-specific alleles were accurately shown to be derived from the lymphocytes and placenta, respectively. Analysis of tissue origin for plasma DNA carrying genetic variants is potentially useful for noninvasive prenatal testing, transplantation monitoring and cancer screening.

## Introduction

The circulation receives DNA from different tissues and organs within the body. The analysis of plasma DNA from specific tissues or organs is useful for revealing and monitoring the pathological processes in different tissues. In scenarios where the genetic composition of a target tissue or organ is different from the host constitutional genome, plasma DNA carrying the tissue- or organ-specific variants can be used to identify DNA molecules released by the tissue or organ. For example, in pregnant women, plasma DNA carrying fetal-specific alleles can be used for prenatal analysis of the fetal genetic constitution (Kitzman et al., 2012; Lo et al., 2010). In organ transplant recipients, the concentrations of donor-specific DNA has been used to reflect the tissue damage associated with acute rejection (De Vlaminck et al., 2014, 2015; Knight et al., 2019; Lo et al., 1998; Schütz et al., 2017). Notably, immediately after organ transplantation, the plasma concentration of donor-derived DNA surges (De Vlaminck et al., 2015). Because of this initial surge, the analysis for donor-derived DNA has limited value in identifying graft rejection and infection during the first 60 days of transplantation (De Vlaminck et al., 2015). The exact mechanism of this initial surge is unclear. It is possible that the hematopoietic cells within the transplanted organ are more likely to release a significant amount of DNA into the circulation during the initial days after the transplantation. However, existing methods for detecting DNA derived from a transplanted organ in plasma rely on identifying genetic differences between the organ donor and the recipient (De Vlaminck et al., 2014, 2015; Knight et al., 2019; Lo et al., 1998; Schütz et al., 2017). These methods cannot be used to further distinguish the exact cell types the donor DNA is derived from.

In situations where the genetic compositions of the different organs are the same, tissue composition analysis based on detecting organ-specific alleles would not be applicable. To overcome this, recent efforts have been made to measure the composition of DNA using epigenetic approaches. These approaches include methylation deconvolution (Moss et al., 2018; Sun et al., 2015), mapping nucleosomal patterns (Snyder et al., 2016; Sun et al., 2019), analysis of end DNA motifs, end positions and jaggedness (Chan et al., 2016; Jiang et al., 2018; Jiang, Sun, et al., 2020; Jiang, Xie, et al., 2020), and the profiling of RNA transcripts (Koh et al., 2014; Tsui et al., 2014). In these methods, the features of interest, e.g. methylation patterns, of the plasma DNA were profiled and compared with those of the candidate tissues. Then the relative contribution of the different tissues to the circulating DNA was determined mathematically. One potential application of plasma DNA tissue composition analysis is to reveal the likely location of a concealed cancer. Recently, it has been shown that the analysis for circulating cell-free tumor DNA (ctDNA) is useful for the screening of early asymptomatic cancers (Chan et al., 2017; Lennon et al., 2020; Liu et al., 2020). As cancer-associated genetic and epigenetic changes are present in virtually all types of cancers (Chan, Jiang, Chan, et al., 2013; Chan, Jiang, Zheng, et al., 2013; Leary et al., 2012; Wong et al., 1999), the detection of these cancer-associated aberrations in plasma can potentially serve as universal tumor markers for the screening of cancers in general. However, how subjects with positive results of a universal cancer test can be further worked up is an important but relatively under-explored topic. In a study by Lennon et al., subjects tested positive with ctDNA test that detected a wide variety of cancers were investigated with whole body positron emission tomography-computed tomography (PET-CT) (Lennon et al., 2020). If the potential tissue origin of the cancer can be obtained from ctDNA analysis, more focused investigations, e.g. high-resolution imaging of an affected organ, can be performed. These organ-specific investigations could provide better sensitivity and specificity and could be achieved with a lower dose of radiation to the patients. In previous proof-of-principle studies, the tissue origin of cancers was successfully revealed by plasma DNA deconvolution (Liu et al., 2020; Moss et al., 2018; Sun et al., 2015). However, existing approaches only allow tissue composition analysis of the whole pool circulating DNA rather than specifically to the tumor-derived DNA. The accuracy of these approaches would be affected by the fractional concentration of tumor-derived DNA in the sample.

In this study, we developed a method called Genetic-Epigenetic Tissue Mapping (GETMap) to determine the tissue composition of plasma DNA carrying genetic variants which are different from the host constitutional genome. This method is based on the comparison of the methylation profiles of the plasma DNA carrying genetic variants and the relevant tissues or organs that plasma DNA is potentially derived from. First, we validated this approach using a pregnancy model through the analysis of the tissue origin of the plasma DNA carrying fetal-specific alleles. Then, we applied this method to measure the tissue compositions of plasma DNA carrying cancer-associated mutations (i.e., present in tumor cells or plasma but absent from buffy coats) in hepatocellular cancer patients and those molecules carrying donor-specific alleles in lung transplant recipients. The former analysis can provide information regarding the tissue origin of the cancer and the latter analysis provided insights on the reason for the surge of donor-derived DNA in the plasma of organ transplant recipients during the early post-transplantation period.

## Results

### Principle of GETMap

The principle of the GETMap analysis is illustrated in Fig. 1. The first step is to identify different sets of plasma DNA molecules based on genotypic differences. For example, the two sets of plasma DNA molecules carrying cancer-associated mutations and wildtype alleles were identified in cancer patients. In organ transplant recipients, three sets of DNA molecules can be identified, including those carrying the host-specific, recipient-specific alleles and alleles shared between the host and recipient. Similarly, three sets of molecules could be identified in the plasma of a pregnant woman, namely those carrying fetal-specific, maternal-specific alleles and alleles shared by the mother and fetus. Then, the tissue compositions were determined for each set of plasma DNA molecules through comparing the methylation profile of the plasma DNA molecules and the methylation profiles of the relevant tissues after bisulfite sequencing. While there are some similarities between the deconvolution step and that described in our previous study (Sun et al., 2015), there are notable differences. First, only DNA molecules of interest, e.g. those carrying fetal-specific alleles, or cancer-associated mutations or donor-derived alleles, are analyzed. Second, only CpG sites near informative SNP alleles are included in the algorithm. The details of the mathematical calculation are described in the Materials and Methods section. For the choice of candidate tissues used for the GETMap analysis, we included the tissues (including neutrophils, lymphocytes, liver, and placenta) that have been validated in a previous study on tissue deconvolution by methylation analysis (Sun et al., 2015). The inclusion of the placenta also allows us to use the analysis of fetal DNA in maternal plasma as a model to validate this new approach. As this study also analyzed patients receiving lung transplantation, lung is further included as one candidate tissue in the plasma DNA deconvolution. The methylation status of the plasma DNA molecules was determined by bisulfite sequencing.

**Fig. 1.**
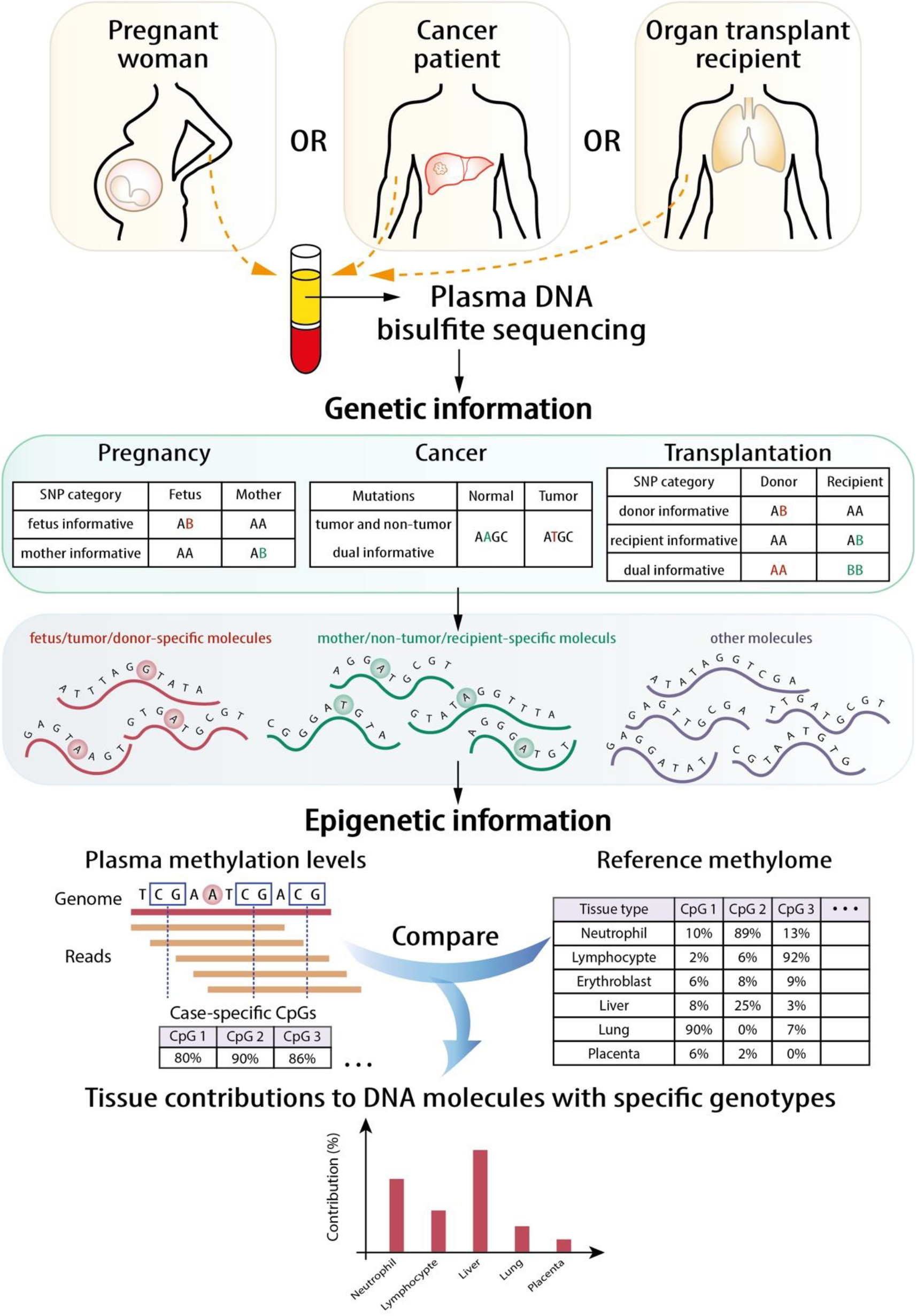
Schematic illustration of the principle of GETMap analysis. The paired individuals (e.g. fetus/mother and organ donor/recipient, and tumor/normal tissue) are genotyped to identify SNP alleles specific for one of them. After bisulfite sequencing, plasma DNA molecules carrying individual-specific alleles and at least one CpG site are identified. The plasma DNA methylome is compared with the methylation profiles of reference tissues to determine the tissue composition of the subset of plasma DNA molecules derived from a particular individual.

### Deconvolution of fetal- and maternal-derived DNA in maternal plasma

We first used the analysis of plasma DNA of pregnant women as a model to demonstrate the feasibility of GETMap. Venous blood samples were collected from 30 pregnant women with 10 in each of the first, second or third trimesters of gestation. Placental tissues were obtained from chorionic villus sampling or amniocentesis for the first and second trimester pregnant women. For third trimester pregnant women, the placenta was collected after delivery. The pregnant woman and the placental tissue were genotyped using the Illumina whole-genome arrays (HumanOmni2.5, Illumina). Based on the genotypes of the mother and fetus, we identified a median of 190,706 (range: 150,168-193,406) maternal-specific informative single nucleotide polymorphisms (SNPs) where the mother was heterozygous and the fetus was homozygous, and a median of 195,331 (range: 146,428-202,800) fetal-specific informative SNPs where the mother was homozygous and the fetus was heterozygous. After bisulfite sequencing of maternal plasma DNA, a median of 103 million uniquely mapped reads (range: 52-186 million) were identified in the maternal plasma DNA samples. Plasma DNA molecules carrying the fetal- and maternal-specific alleles were identified. A median of 162,813 CpG sites (range: 8,237-295,671) and 53,039 CpG sites (range: 16,796-138,284) were identified on the plasma DNA molecules carrying maternal-specific and fetal-specific alleles, respectively. For the plasma DNA molecules carrying fetal-specific alleles, the median deduced contribution from the placenta was 100% (Fig. 2A). These results are compatible to the results of previously studies that fetal DNA in maternal plasma is derived from the placenta (Alberry et al., 2007; Masuzaki, 2004). For molecules carrying maternal-specific alleles, a median of 80% of DNA molecules were deduced to be derived from hematopoietic cells (i.e., neutrophils and lymphocytes) (Fig. 2B). All cases showed no contribution from the placenta. For molecules carrying the shared alleles at SNPs where the mother was homozygous and the fetus was heterozygous, the deduced placental contribution showed a positive correlation with the fetal DNA fractions based on the ratio between the number of plasma DNA molecules carrying fetal-specific alleles and alleles shared by the mother and the fetus (Fig. 2C).

**Fig. 2.**
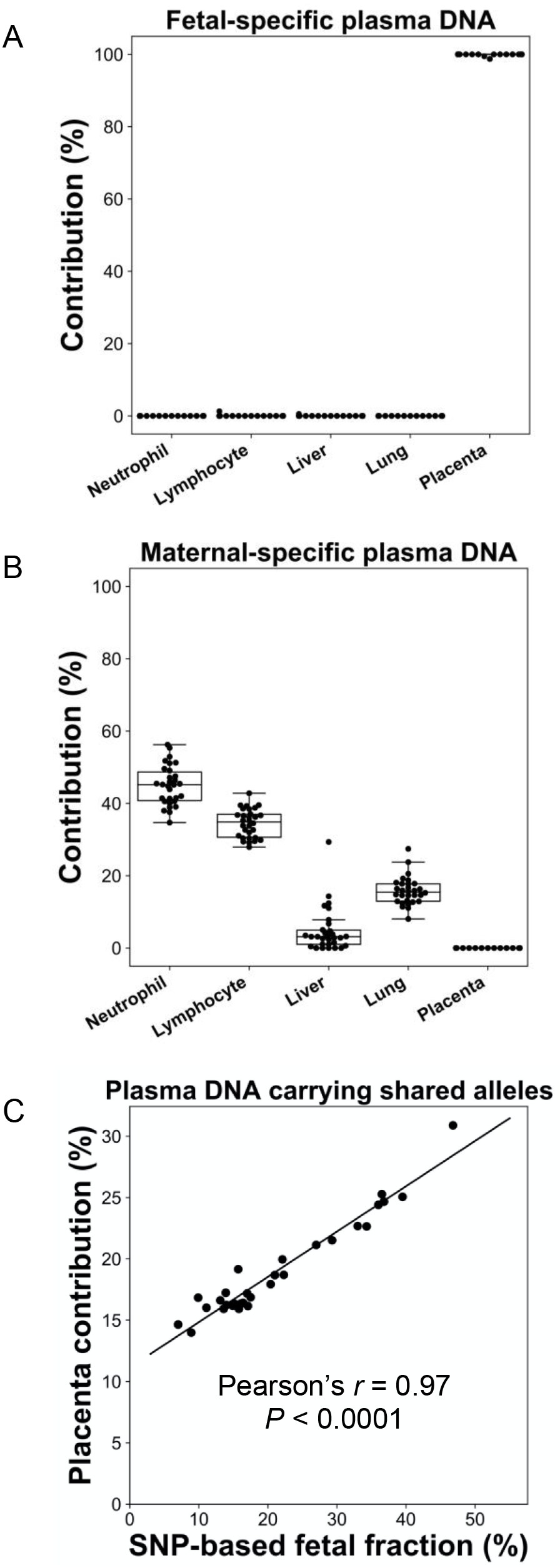
Percentage contributions of different cell types to maternal plasma DNA carrying (A) fetal-specific alleles; and (B) maternal-specific alleles in 30 pregnant women. (C) Correlation between percentage contribution of the placenta to maternal plasma DNA molecules carrying alleles shared by the fetus and mother and SNP-based fetal DNA fraction.

### Deconvolution of DNA molecules carrying donor- and recipient-specific alleles following lung transplantation

We applied GETMap analysis to patients who had received lung transplantation and explored if the tissue composition would change over time. Forty samples from 11 patients were collected (Table S1). By comparing the SNP genotypes between the donor and recipient, we identified a median of 270,144 (range: 254,846-344,024) donor-specific informative SNPs where the donor was heterozygous and the recipient was homozygous and a median of 270,285 (range: 261,529-357,009) recipient-specific informative SNPs where the donor was homozygous and the recipient was heterozygous. In addition, a median of 81,957 (range: 77,196-133,422) dual informative SNPs where both the donor and recipient were homozygous but for different alleles were identified. After bisulfite sequencing of the plasma DNA, a median of 327 million uniquely mapped reads (range: 32-481 million) were obtained for each case. A median of 920,830 (range: 141,065-1,329,292) and 141,794 (range: 12,700-529,211) CpG sites were identified on the plasma DNA molecules carrying recipient- and donor-specific alleles, respectively.

For each subject, the first sample was collected at 72 hours after the transplantation. We performed the GETMap analysis on donor-derived DNA molecules for each sample collected at 72 hours posttransplant (Fig. 3A). The median contribution from the lung to the donor-derived DNA was only 17%. Surprisingly, a substantial proportion of the DNA molecules carrying the donor-specific alleles were contributed from the hematopoietic cells. The median contribution from the neutrophils and lymphocytes combined was 78%. The median deduced contribution from all other tissues was 5% in total.

**Fig. 3.**
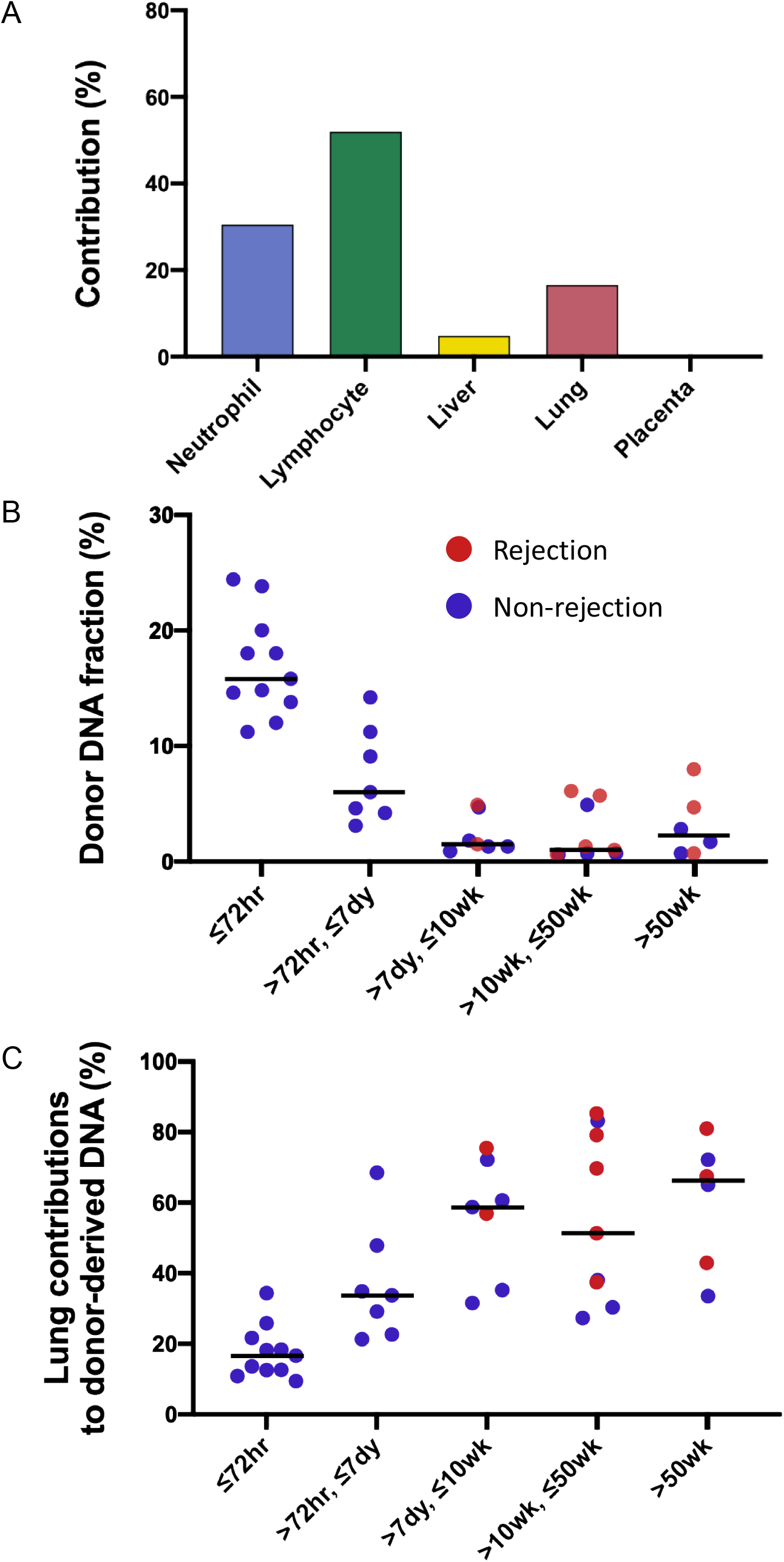
(A) The median percentage contributions of different cell types to plasma DNA carrying donor-specific alleles in patients with lung transplantation at 72 hours posttransplant. (B) Fractional concentrations of donor-derived DNA and (C) percentage contributions of the lung to plasma DNA carrying donor-specific alleles in patients with lung transplantation.

We studied the changes in the lung DNA proportions in the donor-derived plasma DNA molecules with time after transplantation. We categorized the samples based on the time of sample collection posttransplant: within 72 hours; in-between 72 hours, 7 days, 10 weeks, and 50 weeks; and beyond 50 weeks. The 40 samples were thus classified into 5 categories that included 11, 7, 7, 9, and 6 samples, respectively. The median fractional concentrations of donor-derived DNA were 16%, 6%, 2%, 1% and 2% for these categories, respectively (Fig. 3B). The median contributions from the lung to the donor-derived DNA were 17%, 34%, 59%, 51%, 66% for samples in these categories, respectively (Fig. 3C). These data showed that the lung DNA proportions in donor-derived DNA increased with time after transplantation. In contrast, the median contributions from the hematopoietic cells decreased with time, i.e., 78%, 56%, 27%, 41%, and 21% for samples in the 5 categories, respectively. For the plasma DNA molecules carrying the recipient-specific alleles, we observed the hematopoietic cells as the key contributors. For samples in the 5 categories, the median contributions of hematopoietic cells were 83%, 86%, 89 %, 94%, and 84%, respectively (Supplementary Fig. S1).

We further explored if the fractional contribution of the lung to the donor-specific DNA would be useful for the detection of graft rejection. As all the rejection episodes occurred after 7 days, only samples collected after 7 days were used for this analysis. The median donor-derived DNA fractions were 3% for the samples collected during rejection episodes and 1% for those collected during remission (P-value = 0.22, Mann-Whitney rank-sum test, Figure 3B). The median lung contributions were 69% and 48% for these two groups of samples, respectively (P-value = 0.09, Mann-Whitney rank-sum test, Figure 3C).

### Deconvolution of plasma DNA molecules carrying mutant identified in tumor tissues

We then explored if GETMap analysis could reveal the tissue origin of ctDNA in two hepatocellular cancer (HCC) patients. The two patients were denoted as HCC 1 and HCC 2, respectively. In the initial analysis, we first identified the cancer-specific mutations by analyzing the tumor tissues and the buffy coat of the patients. A total of 30,383 and 6,996 tumor-specific single nucleotide mutations were identified from HCC 1 and HCC 2, respectively. After bisulfite sequencing of plasma DNA, 245 and 188 million uniquely mapped reads were obtained for the two patients, respectively. The number of plasma DNA molecules carrying the mutant alleles were 29,868 and 5,090, and these molecules covered 18,193 and 4,076 CpG sites, respectively. Tissue contributions of these tumor-derived plasma DNA molecules were deduced by GETMap analysis (Fig. 4). The liver was deduced to be the key contributor with 90% (HCC 1) and 87% (HCC 2). A small contribution of 10% (HCC 1) and 13% (HCC 2) was from the placenta. The number of molecules carrying the wildtype alleles were 153,238 and 26,792, containing 35,883 and 8,156 CpG sites, respectively. The contribution of the hematopoietic cells was deduced to be 48% (HCC 1) and 53% (HCC 2) whereas the liver contributed 32% (HCC 1) and 23% (HCC 2).

**Fig. 4.**
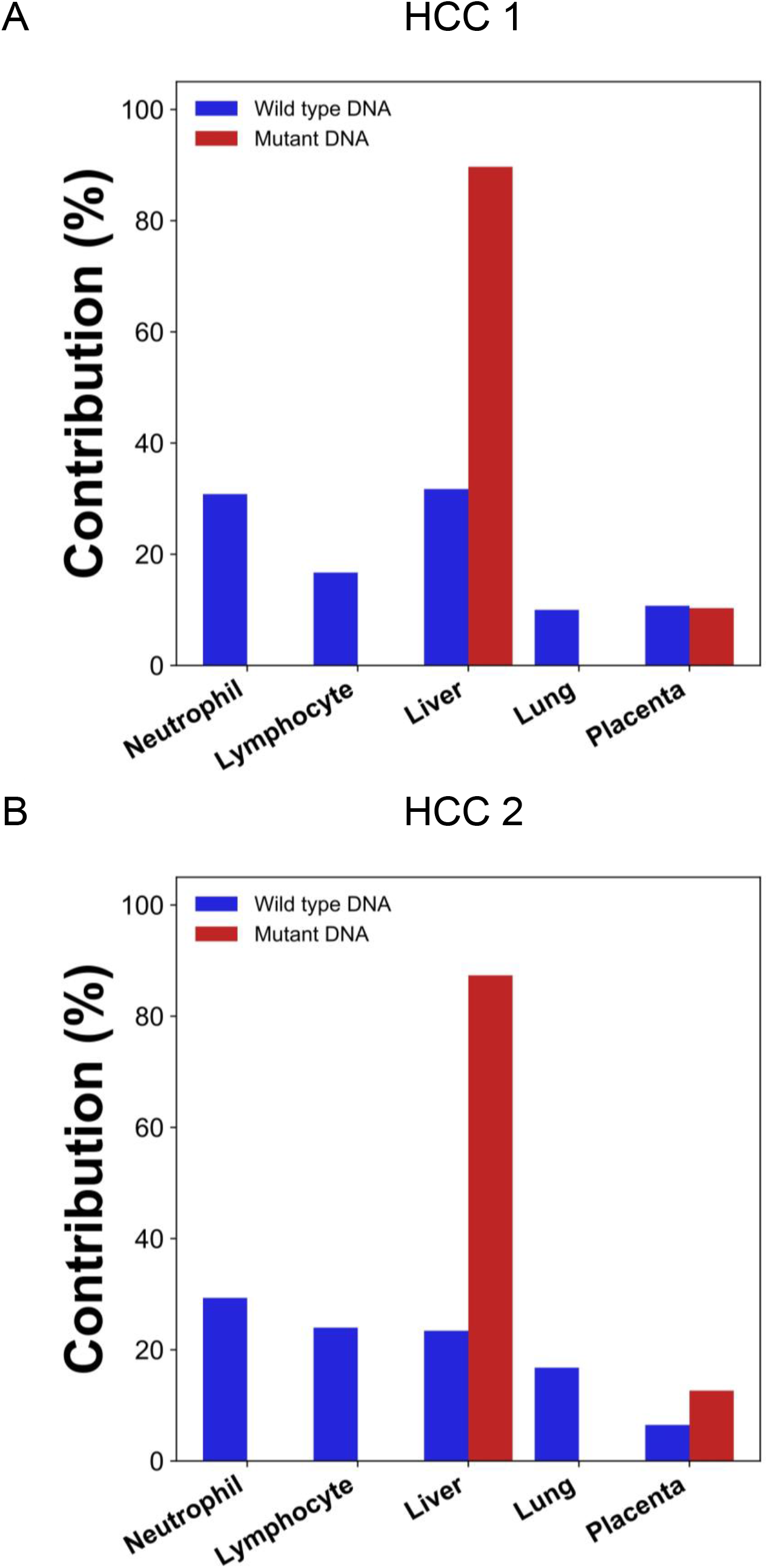
Percentage contributions of different tissues to plasma DNA with tumor-specific and wildtype alleles in two HCC patients. The tumor-specific mutations were deduced from the tumor tissue.

### Deconvolution of DNA carrying mutations directly derived from plasma

In the scenario of cancer screening using a universal tumor marker based on plasma DNA analysis, the tumor tissue would not be available for mutation analysis. Hence, we further explored if the cancer mutations can be directly derived from plasma DNA analysis. To obtain the mutation information directly from the plasma DNA, we sequenced the buffy coat and plasma DNA without bisulfite conversion. The sequencing depth for the plasma DNA were 50× and 61× haploid genome coverage and those for the buffy coat DNA were 53× and 55× in HCC 1 and HCC 2, respectively. Single nucleotides variations present in the plasma for more than a threshold number of occasions but not in the buffy coat were identified as candidate mutations (See details in the Materials and Methods). The number of candidate mutations identified were 10,864 and 3,446 for the two HCC patients. GETMap analysis was then performed using the plasma DNA bisulfite sequencing data. The number of plasma DNA molecules carrying the cancer mutations were 16,200 and 4,112, and covered 12,887 and 2,991 CpG sites, respectively. For molecules carrying mutations, the contributions from the liver were estimated to be 69% (HCC 1) and 95% (HCC 2) (Fig. 5). The placenta contributed the remaining proportion of 31% (HCC 1) and 5% (HCC 2). For molecules carrying wildtype alleles, hematopoietic cells, including neutrophils and lymphocytes, contributed a total of 51% (HCC 1) and 27% (HCC 2).

**Fig. 5.**
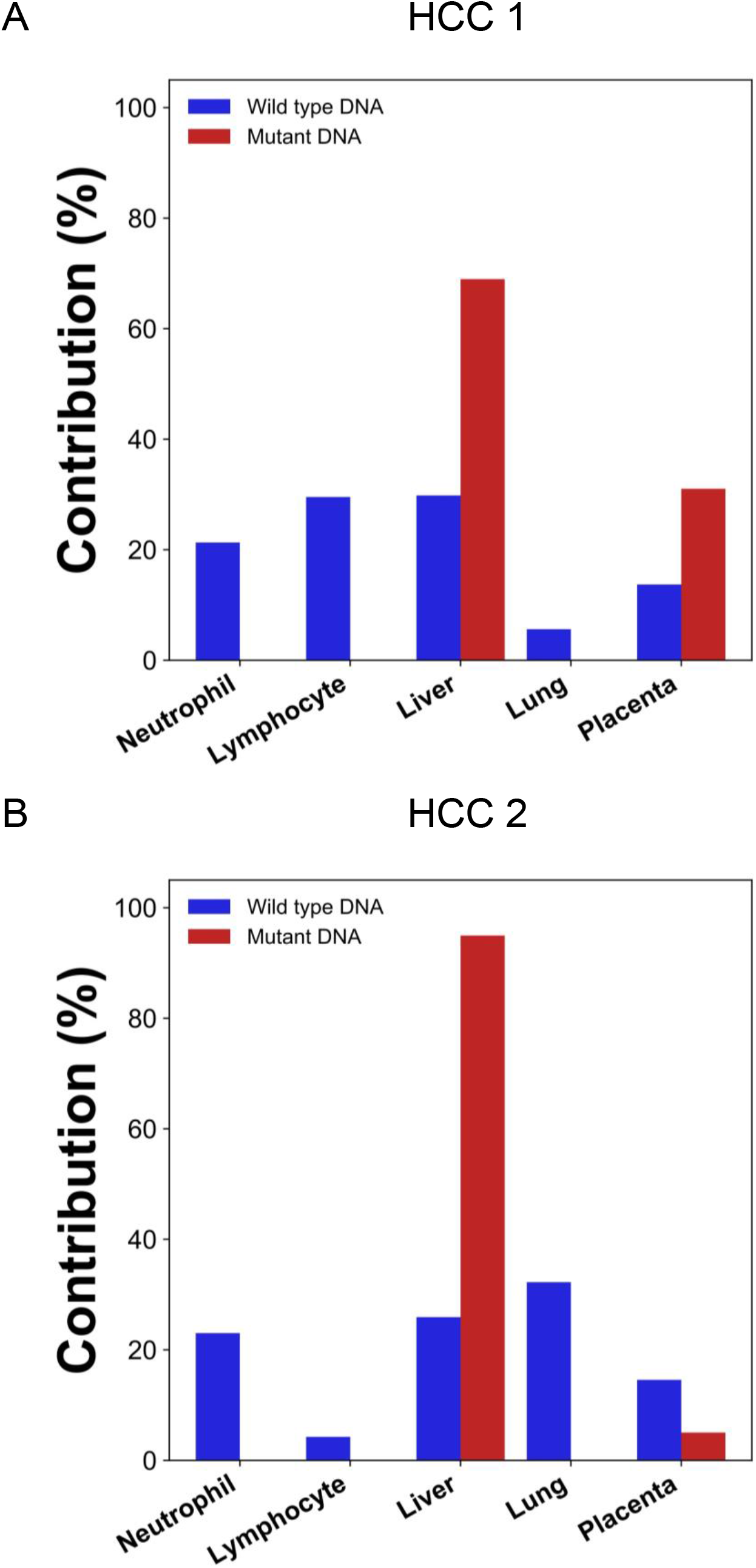
Percentage contributions of different tissues to plasma DNA with tumor-specific and wildtype alleles in two HCC patients. The tumor-specific mutations were deduced directly from the plasma.

### Deconvolution of plasma DNA for a pregnant woman with lymphoma

We previously reported the deconvolution results of total plasma DNA for a pregnant woman who was diagnosed as having follicular lymphoma during early pregnancy (Sun et al., 2015). In the current study, we explored if GETMap analysis could determine the tissue composition of the fetal- and cancer-derived DNA independently. We sequenced the lymphoma tissue, as well as the normal cells harvested from buccal swab and post-treatment buffy coat. As the pregnancy was terminated at time of the diagnosis of cancer, no placental tissue was collected. Hence, we deduced the fetal genotypes directly from the plasma DNA. Based on the non-bisulfite sequencing results of the plasma DNA and normal cells, 254,540 variants were identified in the plasma DNA. The algorithm for classifying these variants into fetal-specific alleles and cancer mutations is shown in Supplementary Fig. S2. We reasoned that variants overlapping with the common variations in the dbSNP Build 135 database were more likely derived from the fetus whereas those not overlapping with the database were more likely to come from the tumor. For the 13,546 variants that did not overlap with dbSNP database, 2,641 were detected in 3 or more sequence reads of the tumor tissues. These variants are regarded as tumor mutations for GETMap analysis. For the 240,994 variants overlapping with the dbSNP database, 231,552 were completely absent in the tumor tissue. These variants were likely derived from the fetus and are regarded as fetal-specific alleles for the GETMap analysis. The allele frequencies for the fetal-specific SNPs and tumor-specific mutations in plasma were normally distributed and peaked at 6% and 20%, respectively (Supplementary Fig. S3).

After bisulfite sequencing of plasma DNA, we obtained 700 million uniquely mapped reads. We identified DNA molecules carrying the tumor-specific mutant alleles, wildtype alleles, fetal-specific alleles, and the alleles shared by the fetus and the mother. The GETMap analysis was performed on each set of plasma DNA molecules to deduce their tissue composition. The number of CpG sites covered by the DNA molecules carrying the mutant and wildtype alleles were 4,781 and 6,660 respectively. For the molecules carrying tumor mutations, it was deduced that 100% was from lymphocytes (Fig. 6A). For molecules carrying the wildtype alleles, the deduced contribution from neutrophils, lymphocytes, liver, lung and placenta were 29%, 46%, 13%, 2% and 11%, respectively. For DNA molecules carrying the fetal-specific, the deduced contribution from the placenta was 95% (Fig. 6B). For those carrying alleles shared by the mother and fetus, the deduced contribution from neutrophils, lymphocytes, liver, lung and placenta were 23%, 48%, 11%, 14% and 5%, respectively.

**Fig. 6.**
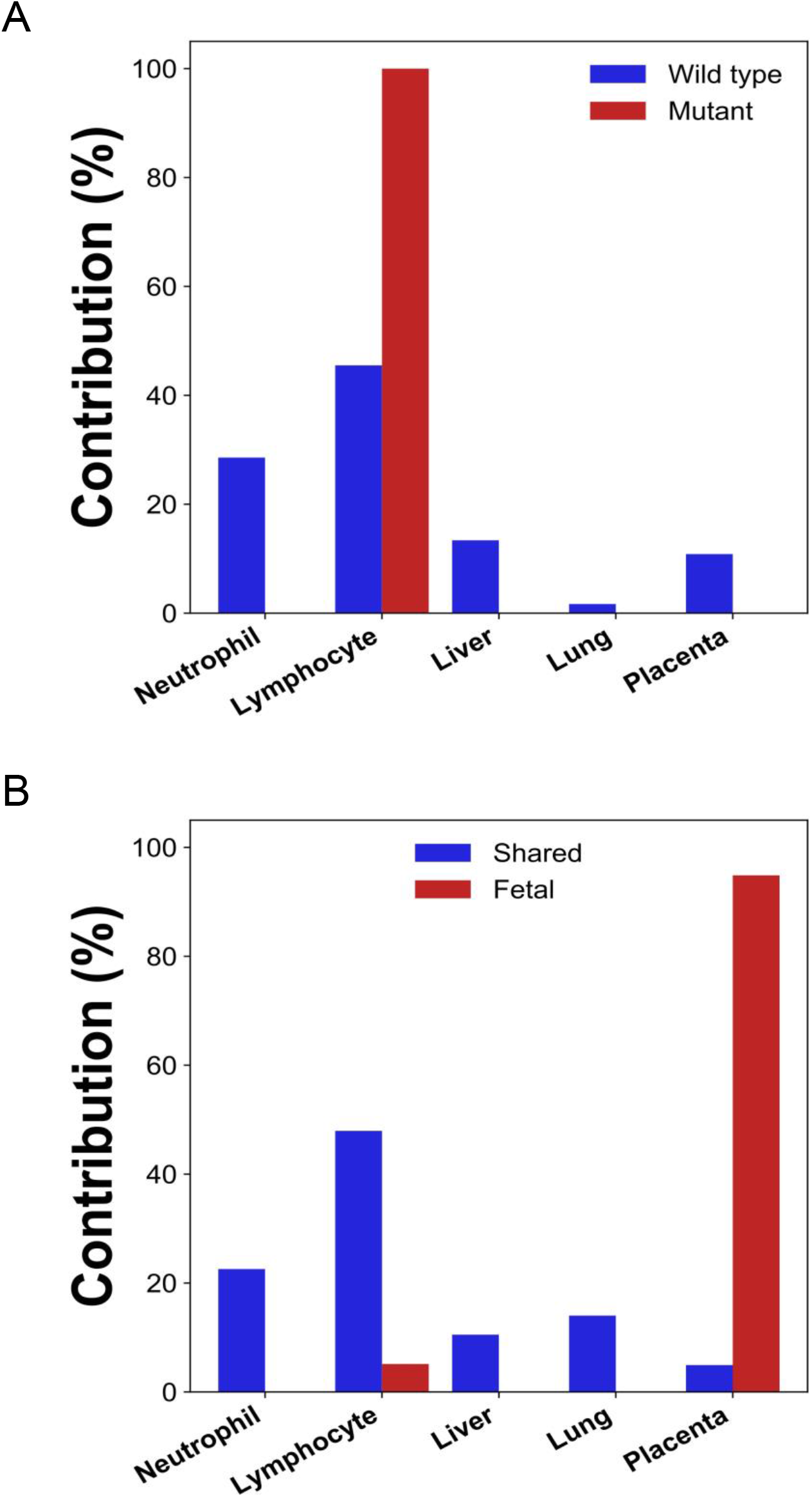
Percentage contributions of different tissues to (A) plasma DNA with tumor-specific and wildtype alleles, and (B) fetal-specific plasma DNA and DNA carrying the alleles shared by the fetus and the mother in a pregnant woman with lymphoma.

## Discussion

In this study, we developed GETMap analysis to determine the tissue origin of plasma DNA molecules carrying genetic variants. In this method, we first identified a subset of plasma DNA molecules carrying specific alleles. Then, by comparing the methylation status of these molecules and the methylation profiles of the candidate tissue organs, we could determine the tissue composition of the DNA molecules. In the first part of the study, we used the pregnancy model to validate the GETMap analysis. The plasma DNA molecules carrying the fetal-specific alleles were deduced to be 100% derived from the placenta. For the molecules carrying the alleles shared by the fetus and the mother, the percentage contribution from the placenta showed a positive linear relationship with the fractional concentration of fetal DNA based on SNP analysis. These results are consistent with the previous studies which showed that the fetal DNA in maternal plasma are indeed derived from the placenta. For the plasma DNA molecules carrying maternal-specific alleles, no contribution from the placenta was observed. A large proportion was derived from the hematopoietic cells, neutrophils and lymphocytes, with a median total contribution of 80%. These figures are comparable to those reported previously in healthy subjects (Gai et al., 2018; Sun et al., 2015). These results demonstrate the feasibility of determining the tissue contributions to the different genetic components of plasma DNA using GETMap analysis.

Then we showed that, in patients who had received lung transplantation, a substantial proportion of donor-derived DNA was derived from the hematopoietic cells during the early post-transplant period. Previous studies have shown that a high level of DNA carrying donor genotypes would be present in the plasma of organ-transplant recipients during the early post-transplant period even in the absence of any evidence of organ rejection (De Vlaminck et al., 2015). Hence, quantitative analysis for donor DNA in plasma cannot be used for reflecting transplant organ damage or rejection within 60 days of transplantation. The reason for this elevation in donor DNA was unclear. Using GETMap analysis, we determined the tissue composition of plasma DNA molecules carrying donor-specific alleles for samples collected at different time intervals after transplantation. Importantly, at 72 hours after transplantation, the median contribution from the lung was only 17% and a substantial contribution of 78% was from hematopoietic cells. This is likely due to the presence of residual blood cells in the transplanted organ and they could release DNA with donor genotypes into the circulation. The contribution of the lung gradually increases with time together with a parallel decline in the contribution of the hematopoietic cells. The median contribution of hematopoietic cells dropped to 21% after 50 weeks. The persistent contribution from the hematopoietic cells may be due to imprecision of measurement as the concentrations of donor DNA after 50 weeks were very low in patients without evidence of rejection. Alternatively, there may be persistence of donor hematopoietic cells in the body of the transplant recipient. In this regard, it has been shown that some immune cells resident in the donor tissue can be long-lived and self-renewing (Gasteiger et al., 2015). The lung fraction appeared to be higher for samples collected during graft rejection compared with those collected during remission. However, the difference did not reach statistical significance. Future studies with larger sample size would be useful to further explore this point.

We then investigated if GETMap analysis could be used to identify the tissue origin of plasma DNA derived from the tumor. Circulating DNA analysis has increasingly been used in the management of cancer patients, in particular for guiding the use of target therapy and monitoring disease progression (Mok et al., 2017; Wan et al., 2020; Yung et al., 2009). Recently, it has been demonstrated that the analysis for cancer-derived DNA in plasma is useful for the screening of cancers in asymptomatic individuals (Chan et al., 2017; Lennon et al., 2020). As genetic and methylation aberrations are present in almost all cancers, the detection of cancer-associated alterations in plasma DNA can potentially serve as a universal tumor marker for a wide variety of cancers. The capability of a tumor marker for picking up multiple types of cancers can greatly enhance the cost-effectiveness of a cancer screening program. However, the lack of tissue or organ specificity of these tests also poses practical challenges on the work up of subjects with positive test results. In the screening study by Lennon et al., subjects tested positive were further investigated with PET-CT to confirm and localize a possible tumor (Lennon et al., 2020). However, if the tissue origin and location can be obtained from the ctDNA analysis, more targeted investigation on the potentially affected organ may be performed. For example, a colonoscopy can be performed for individuals who are suspected of having colorectal cancers. This targeted investigation approach not only provides a more accurate assessment for cancers, it also reduces the radiation exposure of the tested positive subjects. Here, we used the GETMap analysis to determine the tissue origin of plasma DNA carrying cancer-associated mutations. First, we compared the sequencing results of the tumor tissues and the blood cells to identify the mutations in the tumor tissues of two HCC patients. After bisulfite sequencing of the plasma DNA, DNA molecules carrying the cancer-associated mutations were identified and their methylation profiles were used to deduce the contribution from different tissues. The liver was deduced to be the key contributor to these cancer-derived plasma DNA molecules with a contribution of 90% and 87% for the two HCC patients. The remaining portion, i.e. 10% and 13%, were attributed to placental contribution. The attribution of a small proportion of ctDNA to originate from the placenta may be due to the fact that global hypomethylation and hypermethylation of tumor suppressor genes are common features in both the placenta and tumor tissues (Chan, Jiang, Chan, et al., 2013; Feinberg & Vogelstein, 1983; Lun et al., 2013). Although this analysis suggests that GETMap analysis may be useful for revealing the tissue origin of ctDNA, the requirement of tumor tissues for mutation identification limits its practical application in cancer screening. To overcome this, we further attempted to identify cancer mutations directly from plasma DNA sequencing. In this regard, non-bisulfite sequencing for the plasma DNA and the blood cells of the cancer patients were performed. The single nucleotide variants present in the plasma DNA but not in the blood cells were regarded as cancer-associated mutations. GETMap analysis was performed on the plasma DNA molecules carrying these mutations using the bisulfite sequencing data. Despite a smaller number of cancer-associated mutations could be identified by directly sequencing plasma DNA compared with sequencing the tumor tissues, the liver was again correctly identified as the key contributor to these cancer-derived DNA molecules. These results suggest that the GETMap analysis could be useful in revealing the tissue origin and location of a concealed cancer in patients who are screened positive with a tumor marker that detects various types of cancers.

We further challenged GETMap analysis with a complex scenario where a woman developed lymphoma during pregnancy. Her plasma consisted of DNA derived from the lymphoma tissues, the fetus and the normal cells. As fetal tissue was not available, fetal genotypes were deduced by sequencing plasma DNA, maternal blood cells/buccal cells and tumor tissues. Sequence variants present in plasma that overlap with dbSNP database but absent in the tumor tissues were regarded as fetal-specific alleles. Variants detected in plasma and tumor tissues, but not overlap with dbSNP database were regarded as tumor-specific. Plasma DNA molecules carrying these fetal-specific alleles were deduced to be predominantly (95%) derived from the placenta whereas those carrying the tumor-specific alleles were solely from lymphocytes.

There has been increasing interests in the tissue composition circulating cell-free DNA. Methods based on analysis of DNA methylation (Gai et al., 2018; Lehmann-Werman et al., 2016; Sun et al., 2015), nucleosome footprint (Snyder et al., 2016; Sun et al., 2019), sequence motifs, end coordinates and jaggedness (Chan et al., 2016; Jiang et al., 2018; Jiang, Sun, et al., 2020; Jiang, Xie, et al., 2020) have been developed. However, existing methods only allow the deconvolution of all the DNA as a single entity. In contrast, GETMap analysis can determine the tissue origin of subsets of plasma DNA that carry different genetic variations. The specific analysis of a particular component can enhance the signal-to-noise ratio and eliminate the variation caused by the difference in the concentrations of the target DNA, e.g. DNA derived from the tumor. Furthermore, clonal hematopoiesis has been identified as one important source of false-positive results for liquid biopsy-based cancer screening tests. In this regard, GETMap would be useful for identifying the hematopoietic origin of the abnormal signal in such cases. Although the number of cases is relatively small in this proof-of-principle study, we have illustrated the potential applications in cancer detection, prenatal testing and organ transplant monitoring. While the current format of this method is based on whole genome bisulfite sequencing, a targeted sequencing approach enriching for regions with mutation hotspots and differential methylation across different tissues can be developed to enhance the cost-effectiveness of this approach.

## Materials and Methods

### Samples and processing

The project was approved by the Joint Chinese University of Hong Kong-Hospital Authority New Territories East Cluster Clinical Research Ethics Committee. All participants provided written informed consent. Pregnant women and HCC patients were recruited from the Prince of Wales Hospital of Hong Kong. The pregnant woman with lymphoma was recruited from the Hong Kong Sanatorium & Hospital, Hong Kong. Lung transplant recipients were recruited from the National Institutes of Health (NIH). Plasma samples were collected longitudinally at one or several time points after transplantation. Venous blood samples were collected into EDTA-containing tubes and centrifuged at 1,600 *g* for 10 min. The plasma portion was recentrifuged at 16,000 g to remove residual blood cells. DNA from plasma was extracted with the QIAamp Circulating Nucleic Acid Kit (Qiagen).

### Identification of Tumor-specific mutations in HCC patients

We prepared libraries using DNA extracted from the tumor tissue and buffy coat with the TruSeq Nano DNA Library Prep Kit (Illumina). Paired-end (2 × 75 bp) sequencing was performed on the HiSeq4000 system (Illumina). Sequencing data were aligned to the human reference genome using the Burrows-Wheeler Aligner (Li & Durbin, 2010). We compared the data of tumor tissue with that of buffy coat to call the tumor-specific mutations using the Genome Analysis Toolkit (version 4.1.2.0) (McKenna et al., 2010).

To call the tumor-specific mutations directly from the plasma, DNA isolated from the plasma was submitted to library preparation and sequencing. The sequencing data of plasma DNA were then compared with that of the buffy coat to identify the tumor-specific mutations. Single nucleotides variations observed in plasma for more than a threshold number of occasions but not in the buffy coat were identified as candidate mutations. The threshold was based on the total number of sequenced reads covering the variant’s nucleotide position as described in our previous study (Chan et al., 2016). In addition, the sequencing reads covering these candidate mutations were realigned to the reference human genome using a second alignment software which could reduce the number of false-positive results caused by alignment errors as described previously (Chan et al., 2016).

### Identification of Tumor-specific mutations and fetal-specific SNPs in the pregnant women with lymphoma

The DNA extracted from the maternal plasma, tumor cells, and normal cells were summited to library preparation using either the KAPA HTP Library Preparation Kit (Kapa Biosystems) or the TruSeq Nano DNA Library Prep Kit (Illumina) following the manufacturer’s instructions. The 2 × 75 (paired-end mode) cycles of sequencing were performed using the Illumina platforms, including the HiSeq, and NextSeq. To call the plasma-specific variants, we compared the sequencing data of DNA extracted from the maternal plasma with that from the normal cells using the dynamic cutoff algorithm as described previously (Chan et al., 2016). We used the biallelic SNPs downloaded from the dbSNP database (Build 135) to classify the plasma-specific variants. For plasma-specific variants within the dbSNP database, we further filtered out the variants that present in the tumor tissue to obtain the fetal-specific SNPs. For the non-dbSNP variants, the single nucleotide variants observed in at least three molecules from the tumor tissue sequencing data were remained as tumor-specific variants. The bioinformatic pipeline for filtering these mutations was written in Python script.

### Microarray-based genotyping

Pre-transplant blood samples were collected from the donor and recipient. Genomic DNA was extracted from whole blood with the DNeasy Blood and Tissue Kit (Qiagen) and amplified with REPLI-g Mini Kit (Qiagen). For the pregnant case, genomic DNA of the mother and fetus were extracted from maternal buffy coat and fetal placenta tissue with the QIAamp DNA Mini Kit (Qiagen). Genotyping was performed on Illumina whole-genome arrays (HumanOmni2.5 or HumanOmni1) following the manufacturer’s protocol (De Vlaminck et al., 2014).

### Bisulfite-treated DNA libraries preparation and sequencing analysis

Libraries were prepared from plasma DNA with the TruSeq Nano DNA Library Prep Kit (Illumina). DNA libraries were subjected to two rounds of bisulfite modification with the EpiTect Bisulfite Kit (Qiagen) following by 12-cycles of PCR amplification. Bisulfite-treated libraries were sequenced in paired-end mode (2 × 75 bp) on a HiSeq 4000 system (Illumina). The sequencing reads were trimmed to remove adapter sequences and low-quality bases (i.e., quality score < 5). The trimmed reads were aligned to the human reference genome build hg19 with Methy-Pipe (Jiang et al., 2014).

### GETMap analysis

We analyzed whole-genome bisulfite sequencing data of different tissues or cell types from public datasets, including BLUEPRINT Project (Martens & Stunnenberg, 2013), Roadmap Epigenomics (Roadmap Epigenomics Consortium et al., 2015), ENCODE (Davis et al., 2018) and GEO (Barrett et al., 2012) and methylomes generated by our group. The methylation levels for 28,217,006 CpG sites across 6 types of tissues were determined. We retrieved the methylation levels of different tissues across the set of CpG sites covered by donor- and recipient-specific DNA molecules. The measured CpG methylation levels of donor- or recipient-specific DNA molecules were recorded in a vector (X) and the retrieved reference methylation levels across different tissues were recorded in a matrix (M). The proportional contributions (P) from different tissues to donor- or recipient-specific DNA molecules were deduced by quadratic programming:

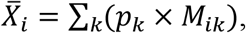

where 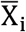 represents the methylation density of a CpG site i in the DNA mixture; pk represents the proportional contribution of cell type k to the DNA mixture; M_ik_ represents the methylation density of the CpG site i in the cell type k. When the number of sites is the same or larger than the number of organs, the values of individual p_k_ could be determined. To improve the informativeness, the CpG sites showed small variability of methylation levels across all reference tissue types were discarded. The remaining CpG sites were characterized with the coefficient of variation (CV) of methylation levels across different tissues less than 0.3 and a difference between a maximum and a minimum methylation levels among tissues more than 25%.

Additional criteria were included in the algorithm to improve the accuracy. The aggregated contribution of all cell types would be constrained to be 100%:

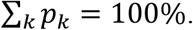

Furthermore, all the organs’ contributions would be required to be non-negative:

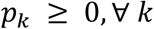

The GETMap deconvolution analysis was performed with a program written in Python (http://www.python.org/).

## Acknowledgement

This work was supported by the Research Grants Council of the Hong Kong SAR Government under the Theme-based research scheme (T12-403/15-N and T12-401/16-W), a collaborative research agreement from Grail and the Vice Chancellor’s One-Off Discretionary Fund of The Chinese University of Hong Kong (VCF2014021). Y.M.D. Lo is supported by an endowed chair from the Li Ka Shing Foundation.

## Supplementary figures and tables

**Fig. S1.**
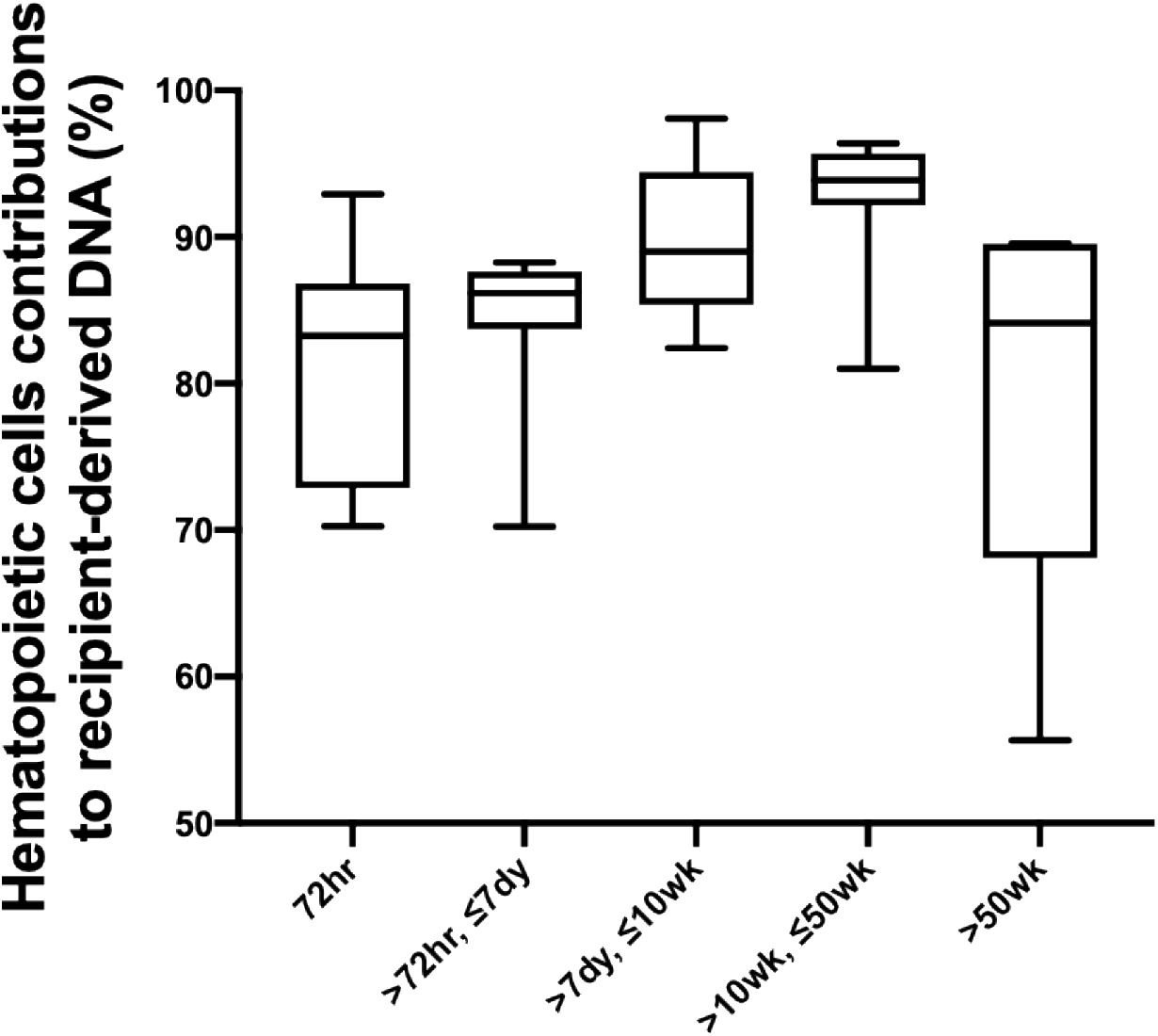
Percentage contributions of hematopoietic cells to the plasma DNA carrying recipient-specific alleles in patients with lung transplantation.

**Fig. S2.**
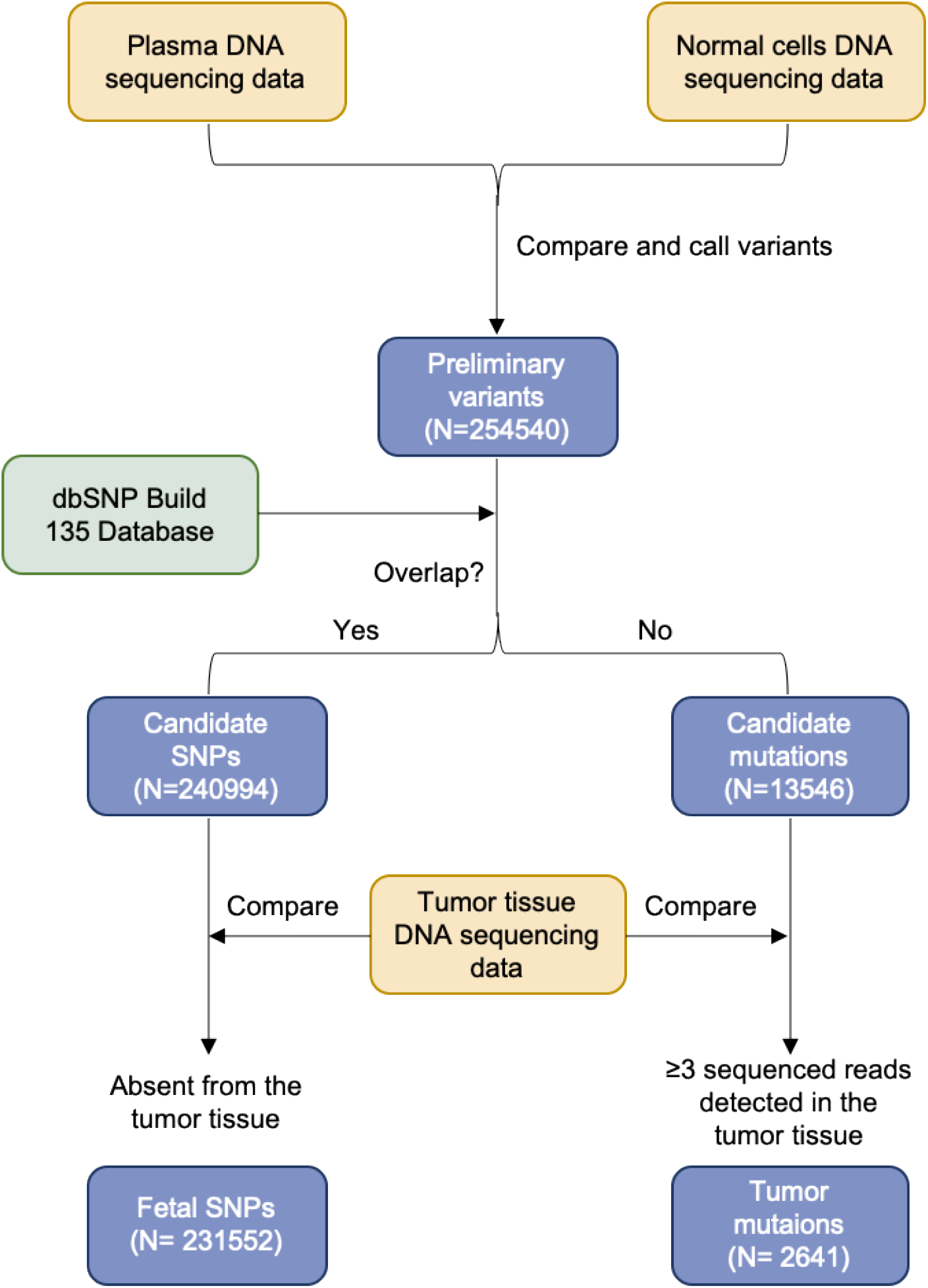
Flowchart of the steps for identifying the fetal-specific alleles and cancer mutations in the pregnant woman with lymphoma.

**Fig. S3.**
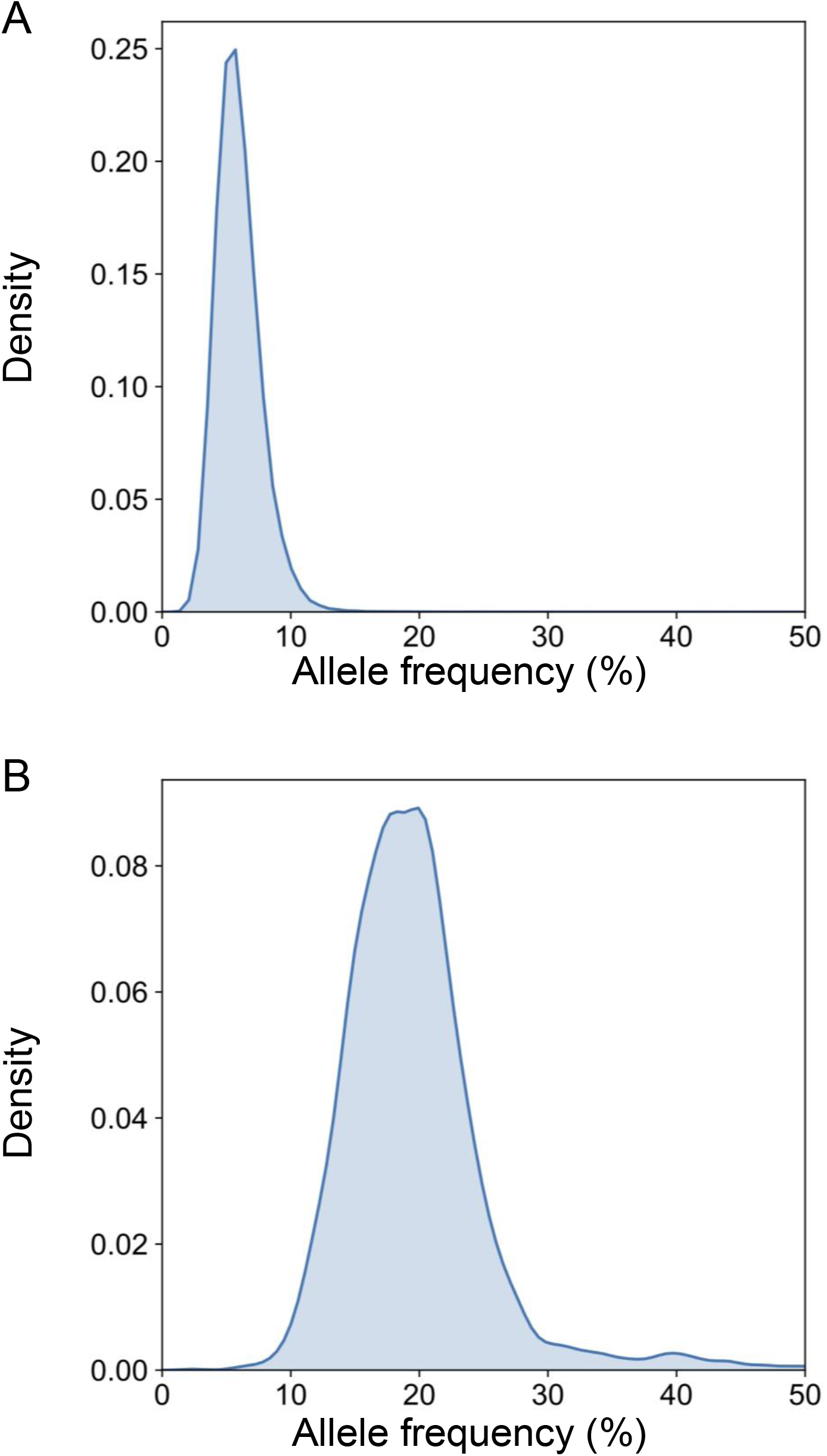
The distribution of the allele frequency of (A) the fetal-specific alleles and (B) the mutant alleles in the plasma of the pregnant woman with lymphoma.

**Table S1.** The demographic profiles of lung transplant recipients.

| Case number | Recipient age | Recipient gender | Donor age | Donor gender | Diagnosis for transplant | Single/Double lung | Cause of death | Time of sample collection posttransplant |
| --- | --- | --- | --- | --- | --- | --- | --- | --- |
| 1 | 34 | M | 32 | M | Cystic fibrosis | double | alive | 72hr |
| 2 | 59 | F | 27 | F | Interstitial lung disease | double | alive | 72hr |
| 3 | 53 | M | 20 | M | Interstitial lung disease | double | alive | 72hr |
| 4 | 63 | M | 16 | F | Interstitial lung disease | double | alive | 72hr, 6dy |
| 5 | 55 | F | 36 | F | Interstitial lung disease | double | alive | 72hr, 7dy |
| 6 | 66 | M | 48 | F | Interstitial lung disease | single | alive | 72hr, 4wk |
| 7 | 66 | F | 18 | M | Chronic obstructive pulmonary disease | single | alive | 72hr, 7dy, <u>5wk</u>, <u>20wk</u>, <u>25wk</u>, <u>157wk</u> |
| 8 | 32 | F | 39 | M | Cystic fibrosis | double | alive | 72hr, 7dy, <u>8wk</u>, 38wk, <u>77wk</u>, 129wk |
| 9 | 67 | F | 53 | F | Sarcoidosis | double | respiratory failure | 72hr, 7dy, 6wk, 13wk, <u>22wk</u> |
| 10 | 44 | M | 35 | F | Re-transplant | double | alive | 72hr, 7dy, 10dy, 4wk, 14wk, <u>25wk</u>, 103wk |
| 11 | 67 | F | 32 | M | Pulmonary arterial hypertension | single | alive | 72hr, 7dy, 5wk, 15wk, <u>26wk</u>, <u>61wk</u>, 104wk |

\* Samples collected when the patient was having a rejection episode were underlined.

